# Robust neural tracking of linguistic speech representations using a convolutional neural network

**DOI:** 10.1101/2023.03.30.534911

**Authors:** Corentin Puffay, Jonas Vanthornhout, Marlies Gillis, Bernd Accou, Hugo Van hamme, Tom Francart

## Abstract

**Objective:** When listening to continuous speech, populations of neurons in the brain track different features of the signal. Neural tracking can be measured by relating the electroencephalography (EEG) and the speech signal. Recent studies have shown a significant contribution of linguistic features over acoustic neural tracking using linear models. However, linear models cannot model the nonlinear dynamics of the brain. To overcome this, we use a convolutional neural network (CNN) that relates EEG to linguistic features using phoneme or word onsets as a control and has the capacity to model non-linear relations.

**Approach:** We integrate phoneme- and word-based linguistic features (phoneme surprisal, cohort entropy, word surprisal and word frequency) in our nonlinear CNN model and investigate if they carry additional information on top of lexical features (phoneme and word onsets). We then compare the performance of our nonlinear CNN with that of a linear encoder and a linearized CNN.

**Main results:** For the non-linear CNN, we found a significant contribution of cohort entropy over phoneme onsets and of word surprisal and word frequency over word onsets. Moreover, the non-linear CNN outperformed the linear baselines.

**Significance:** Measuring coding of linguistic features in the brain is important for auditory neuroscience research and applications that involve objectively measuring speech understanding. With linear models, this is measurable, but the effects are very small. The proposed non-linear CNN model yields larger differences between linguistic and lexical models and, therefore, could show effects that would otherwise be unmeasurable and may, in the future, lead to improved within-subject measures and shorter recordings.

**Index Terms:** EEG decoding, speech processing, CNN, linguistics.

## 1. Introduction

When someone listens to sounds, the information picked up by the ears is carried along the auditory pathway from the cochlea to the brain. The resulting brain response can be measured using a non-invasive method called electroencephalography (EEG). In early studies (e.g., Picton et al., 2005; Anderson et al., 2013), unnatural periodic stimuli are presented to listeners, and the recorded EEG signal is averaged to obtain the resulting brain response, and to enhance its speech-related component.

These short and repeated stimuli do not reflect real-life situations where humans listen to continuous speech that is not repeated. To investigate continuous speech processing, a method that models the mapping between the speech features and the brain response was implemented and described in prior work (Ding and Simon, 2012; Crosse et al., 2016). Such models are linear regressions that either aim to predict the EEG signal from the speech features (i.e., forward modeling), or to reconstruct speech features from the EEG (i.e., backward modeling). Typically, the correlation between the predicted (or reconstructed) EEG (or speech features) and the ground truth is computed to measure what is referred to as neural tracking of speech (Ding and Simon, 2012; Crosse et al., 2016; Vanthornhout et al., 2018; Lesenfants et al., 2019; Brodbeck and Simon, 2020).

Other paradigms to measure neural tracking have been used, such as match-mismatch (MM) classification tasks (de Cheveigńe et al., 2021), which consist of classifying whether a given EEG segment is synchronized (matched) or not with a given speech feature segment. The accuracy obtained on the MM task can be used as a measure of neural tracking.

The speech signal contains multiple features known to be processed at different stages along the auditory pathway. The neural tracking of different features of speech has hence been investigated including:

- Acoustics (e.g., spectrogram, speech envelope (Di Liberto et al., 2015), f0 (Puffay et al., 2022))
- Lexical features (e.g., phoneme onsets, word onsets, (Di Liberto et al., 2015; Lesenfants et al., 2019))
- Linguistics (e.g., phoneme surprisal, word frequency, (Gillis et al., 2022; Brodbeck et al., 2018; Broderick et al., 2018; Weissbart et al., 2019; Koskinen et al., 2020))

Linguistic features are related to the information carried by a word or a phoneme; as such, their presence in the EEG can indicate speech understanding (Gillis et al., 2022, e.g.,). However, different speech features are derived from the same acoustic signal and are often highly correlated (Daube et al. (2019)). Hence, some studies (Gillis et al. (2022); Brodbeck et al. (2018)) control for acoustics and lexical information by measuring the added value of linguistics over acoustics and phoneme/word onsets, effectively comparing a control model with acoustic and lexical features, to a model that also includes linguistic features. This approach is very conservative as the information that is shared between acoustic, lexical, and linguistic features is lumped into the control model. The added value of linguistic features on the model’s performance thus only measures their unique contribution. On the other hand, some studies do not control for any speech feature (e.g., Broderick et al. (2018); Weissbart et al. (2019)), which does not guarantee that the neural tracking measured is not purely acoustic.

Considering the nonlinear response of the brain, and non-stationarities, for instance, due to attentional modulations, the use of linear models is a crude simplification. Following-up recent advances in deep learning in Automatic Speech Recognition (ASR), many studies attempted to relate EEG to speech features using deep learning models (e.g., de Taillez et al. (2020); Thornton et al. (2022); Accou et al. (2021b,a); Monesi et al. (2020); Puffay et al. (2022)); for a review on deep learning models, see Puffay et al. (2023). It is crucial to observe that in these deep learning studies, acoustic features were predominantly employed, while linguistic features were never considered.

Regarding deep-learning-based models, a recent study (Puffay et al. (2022)) implemented a multi-input feature convolutional neural network, which enables investigating the added value of one speech feature over another. Such architecture could enable controlling that the quantified linguistic neural tracking has a component unrelated to e.g., lexical feature.

In this article, we attempt to (1) Measure neural tracking of linguistic features that is unrelated to neural tracking of onsets; and (2) Quantify the impact of deep neural networks over linear models on this task.

In Section 2, we first introduce the speech features we use: lexical features (i.e., phoneme and word onsets) and linguistic features. Lexical features are binary (for each time sample, gets the value of 1 if a word or a phoneme onset occurs, 0 otherwise), and linguistic features modulate the magnitude of non-zero values from word or phoneme onset features with the amount of linguistic information. We then define the MM task that will be used to objectively measure neural tracking, the architectures we used, and the corresponding training paradigm. Lastly, we define the role of lexical control models in measuring the neural tracking of linguistic features while considering their correlation to onsets. To fulfill our two objectives, we split our work into two experiments and report our results in Section 3. First, we investigate the potential added value of our linguistics models over lexical control models at both the phoneme and word levels. Second, we quantify the added value of linguistic features across architectures by comparing the difference in accuracy between linguistics and control models, which is indicative of the power of the model to detect the coding of linguistic features in the EEG.

## 2. Methods

### 2.1. Feature extraction and pre-processing

#### 2.1.1. *Speech features* This paper addresses decoding 6 features from EEG signals. These features are as follows

Phoneme-level lexical feature: onset of any phoneme (PO)
Phoneme-level linguistic features: cohort entropy (CE), and phoneme surprisal (PS)
Word-level lexical feature: onset of any word (WO)
Word-level linguistic features: word frequency (WF), and word surprisal (WS)

Among other linguistic features, we selected those which showed an added value over control speech features in a previous linear model study (Gillis et al., 2022).

### Phoneme onsets and word onsets

Time-aligned sequences of phonemes and words were extracted by performing a forced alignment of the identified phonemes (Duchateau et al., 2009). The resulting features were one-dimensional arrays with pulses on the onsets of, respectively, phonemes and words. Silence onsets were set to 0 for both phonemes and words.

### Active cohort of words

Prior to introducing phoneme-based linguistics, the active cohort of words must be defined. Following previous studies’ definition (Brodbeck et al., 2018; Gillis et al., 2022), it is a set of words that starts with the same acoustic input at any point in the word. Should we find cohorts in English, the active cohort of words for the phoneme */n/* in “ban” corresponds to the ensemble of words that exist in that language starting with “ban” (e.g., “banned”, “banana”, “bandwidth” etc.). For each phoneme, the active cohort was determined by taking word segments that started with the same phoneme sequence from the lexicon.

### Lexicon

The lexicon for determining the active cohort was based on a custom pronunciation dictionary maintained at our laboratory (created manually and using grapheme-to-phoneme conversion; containing 9157 words). As some linguistic features are based on the word frequency in Dutch, the prior probability for each word was computed, based on its frequency in the SUBTLEX-NL database (Keuleers et al., 2010).

### Phoneme-based linguistics

Two phoneme-based linguistic features were extracted from the speech: PS and CE as defined in previous studies (Brodbeck et al. (2018); Gillis et al. (2022)). The two features are derived from the active cohort of words.

PS is a measure of how surprising a phoneme is considering the active cohort and is defined in Equation 1. PS of a given phoneme is defined as the negative logarithm of the conditional probability of each phoneme given the preceding phonemes in the same word. *i* is the phoneme index in the given word. For *PS*_1_, the surprisal corresponding to the first phoneme of the word, we took the frequency of this phoneme at the beginning of a word in the corpus.

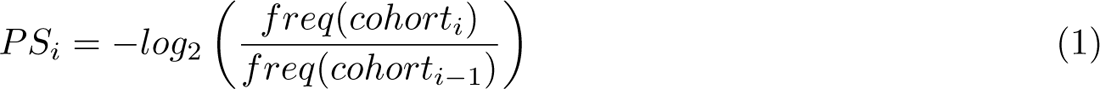

CE reflects the degree of competition among possible words that can be created from the active cohort including the current phoneme. It is defined as the Shannon entropy of the active cohort of words at each phoneme as explained in Brodbeck et al. (2018) (see Equation 2). *CE_i_*is the entropy at phoneme *i* and *p_word_* is the probability of the given word in the language. The sum iterates over words from the active cohort *cohort_i_*. Phoneme-level features are illustrated in Figure 1 (right panel).

**Figure 1:**
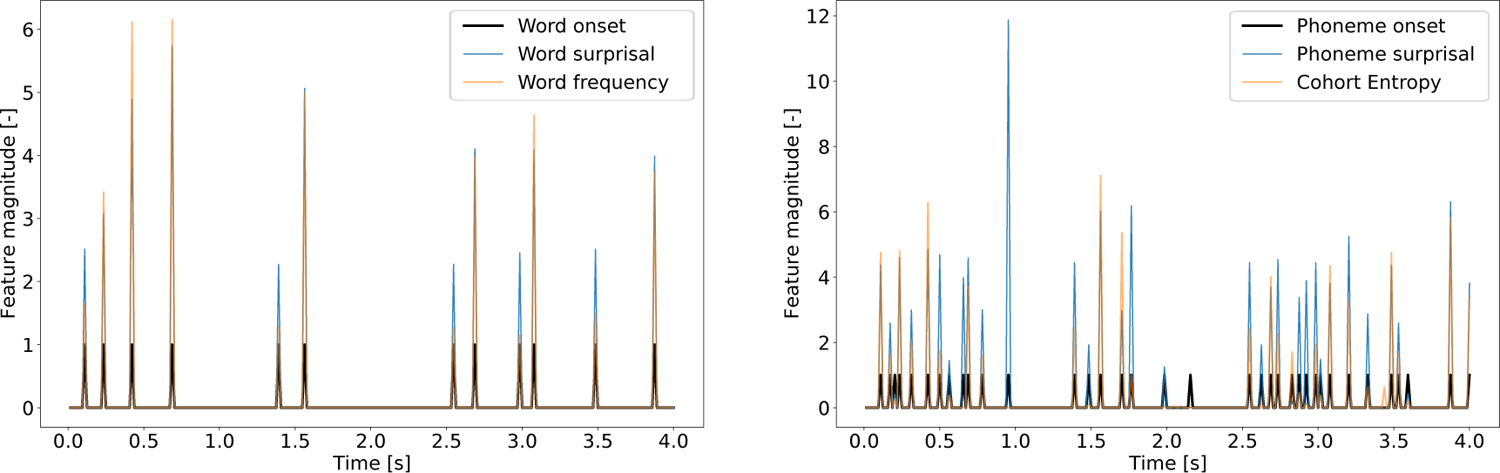
Example word- and phoneme-level lexical and linguistic features.

### Word-based linguistics

Two word-based linguistic features were extracted from the speech: word frequency (WF) and word surprisal (WS) as defined in previous studies (Brodbeck et al. (2018); Gillis et al. (2022)). We used a five-gram with a vocabulary of 400,000 words, trained with Kneser-Ney smoothing (Kneser and Ney, 1995) on newspapers and magazines totaling over 3 billion words *‡*. The prior probability for each word was based on its frequency in the SUBTLEX-NL database (Keuleers et al., 2010). Values corresponding to words not found in the SUBTLEX-NL nor by the five-gram model were set to 0.

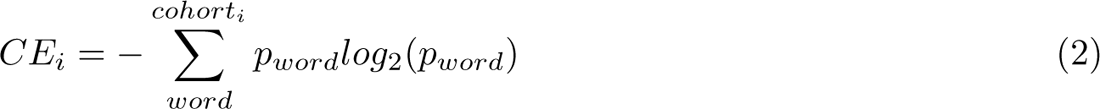

*‡* More recent open-source models such as LlaMa or GPT2 would possibly give a better model of word probabilities.

WF is a measure of how frequently a word occurs in the language, and is defined in Equation 3. WS reflects how surprising a word *w_i_* is considering the four preceding words, as defined in Equation 4. *i* is the index of a given word.

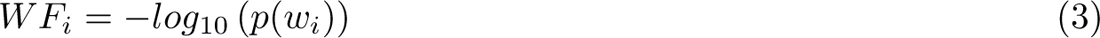

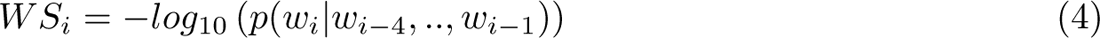

Word-level features are illustrated in Figure 1 (left panel).

#### 2.1.2. Pre-processing

The EEG signal was first downsampled to 1024 Hz. A multi-channel Wiener filter (Somers et al., 2018) was then used to remove eyeblink artifacts, and re-referencing was performed to the average of all electrodes. The resulting signal was band-pass filtered between 1 and 32 Hz using a zero-phase Chebyshev type-II filter with 80 dB attenuation at 10% outside the pass-band and a pass-band ripple of 1 dB, then downsampled to 64 Hz.

As lexical and linguistic features are discrete, they were used as-is. We calculate the features at the 64 Hz sampling rate.

Furthermore, we divided each subject’s EEG and speech features into training, validation, and testing sets as in previous studies using the same stimuli (Monesi et al. (2020); Bollens et al. (2022); Accou et al. (2021b)). We used the validation set for regularization (i.e., early stopping on the validation loss with a patience of 5). The first and last 40% of the recording segment were used for training. The first half of the remaining 20% was used for validation and the last half for testing. For the training, validation, and testing set, the mean EEG signal per channel and corresponding standard deviation were computed. For each set (training, validation, and testing, respectively), each EEG channel was then normalized by subtracting the mean and dividing by the standard deviation previously computed.

### 2.2. Match-mismatch classification task

In this study, we use the performance of the MM classification task to measure the neural tracking of different speech features. This paradigm is depicted in Figure 2.

**Figure 2:**
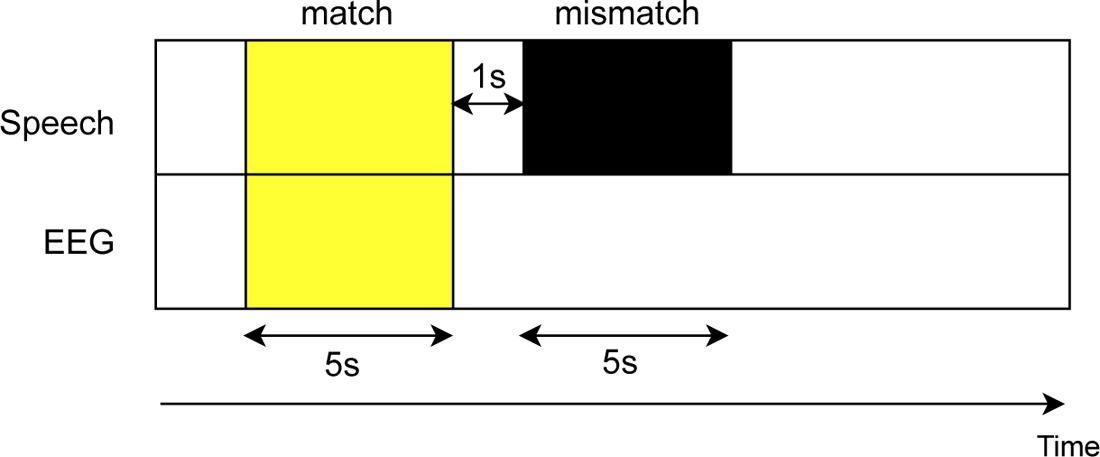
Match-mismatch classification task. The match-mismatch task is a binary classification paradigm that associates the herewith yellow EEG and speech segments. The matched speech segment is synchronized with the EEG (yellow segment), while the mismatched speech occurs 1 second after the end of the matched segment (black segment). The figure depicts segments of 5 s; however, different durations are picked throughout our analyses.

The model is trained to associate the EEG segment with the matched speech segment among two presented speech segments. The matched speech segment is synchronized with the EEG, while the mismatched speech segment occurs 1 second after the end of the matched segment. These segments are of fixed length, namely 10 s for word-based features and 5 s for phoneme-based features, to provide enough context to the models. This task is supervised as the matched, and mismatched segments are labeled. The evaluation metric is classification accuracy.

### 2.3. Model architectures

#### 2.3.1. Preamble

All the described models were created in Tensorflow (2.3.0) (Abadi et al., 2015) with the Keras API ((Chollet et al., 2015))

In this section, we present a linear encoder baseline (Section 2.3.2), a nonlinear CNN (Section 2.3.3), and a linearized CNN (Section 2.3.4) to investigate whether word- and phoneme-based linguistic features carry non-redundant information over word and phoneme onsets respectively.

We then define in Section 2.5 the lexical control models (i.e., without linguistics information) and the linguistics models. In addition, we motivate the use of lexical control models to measure the neural tracking of linguistics.

Finally, we introduce in Section 2.6 the statistical models used to compare linguistic and control model performances across various model architectures and training paradigms; and we briefly explain our data collection procedure in Section 2.7.

#### 2.3.2. Linear encoder baseline

To relate to the existing literature, we evaluated a linear architecture for the lexical control and the linguistics models. In this case, neural tracking is generally measured by computing the correlation between the predicted and ground truth EEG signal. To obtain a performance on the MM task, we implement a correlation-based MM task as depicted in Figure 3.

**Figure 3:**
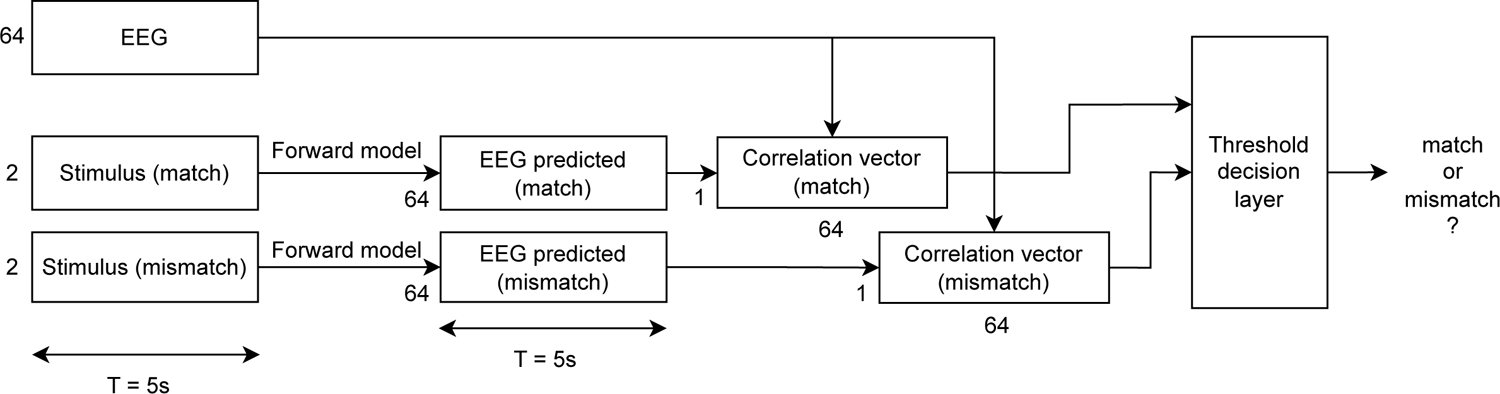
Linear encoder baseline architecture. This model integrates a pre-trained forward model in a correlation-based MM task to estimate the match-mismatch accuracy obtained by a state-of-the-art linear encoder.

We first trained the linear encoder (i.e., a forward model) using an iterative approach, like for a deep neural network (DNN), to maximize the similarity. The forward model applies a linear convolution (kernel size: 600 ms, 64 spatial filters, causal padding) to two concatenated speech features provided as inputs (batch’s dimensions: 2 *× T*, *T* = 30 *s*). We used the mean-squared error (MSE) between the predicted and ground truth EEG as the loss function. We selected the Adam optimizer with a learning rate of 0.001. We also verified that the MSE values reached were in the order of those obtained with the closed-form solution approach in prior studies.

To enable performance comparison with the linearized and nonlinear CNNs, we integrated the pre-trained forward model in the MM task, as depicted in Figure 3. An EEG segment and two speech segments (matched and mismatched) of length *T* = 5 *s* were provided as inputs. The EEG segment has dimensions 64 *× T* and the matched and mismatched speech segments have dimensions 2 *× T* as 2 speech features were systematically concatenated (i.e., either two lexical features for control models or a lexical and a linguistic feature for linguistic models).

The pre-trained forward model was first applied to the matched and mismatched speech segments, resulting in two predicted EEG segments of dimensions 64 *× T*. The Pearson correlation was then computed between corresponding channels of matched and mismatched segments and the ground truth EEG segment, which resulted in two correlation vectors (dimensions: 1 *×* 64). The threshold decision layer counts the number of channels and performs a majority vote. The accuracy was computed from the number of correctly classified segments and computed for each subject.

#### 2.3.3. Nonlinear CNN The nonlinear

CNN was first introduced for this particular task by Puffay et al. (2022) and inspired by Accou et al. (2021b). It enables providing information from *N* different speech features (here *N* = 2) to our model and evaluating its performance on the MM task. By combining multiple features to solve the same MM task, we can investigate whether the model benefits from adding some features or not. The model is depicted in Figure 4.

**Figure 4:**
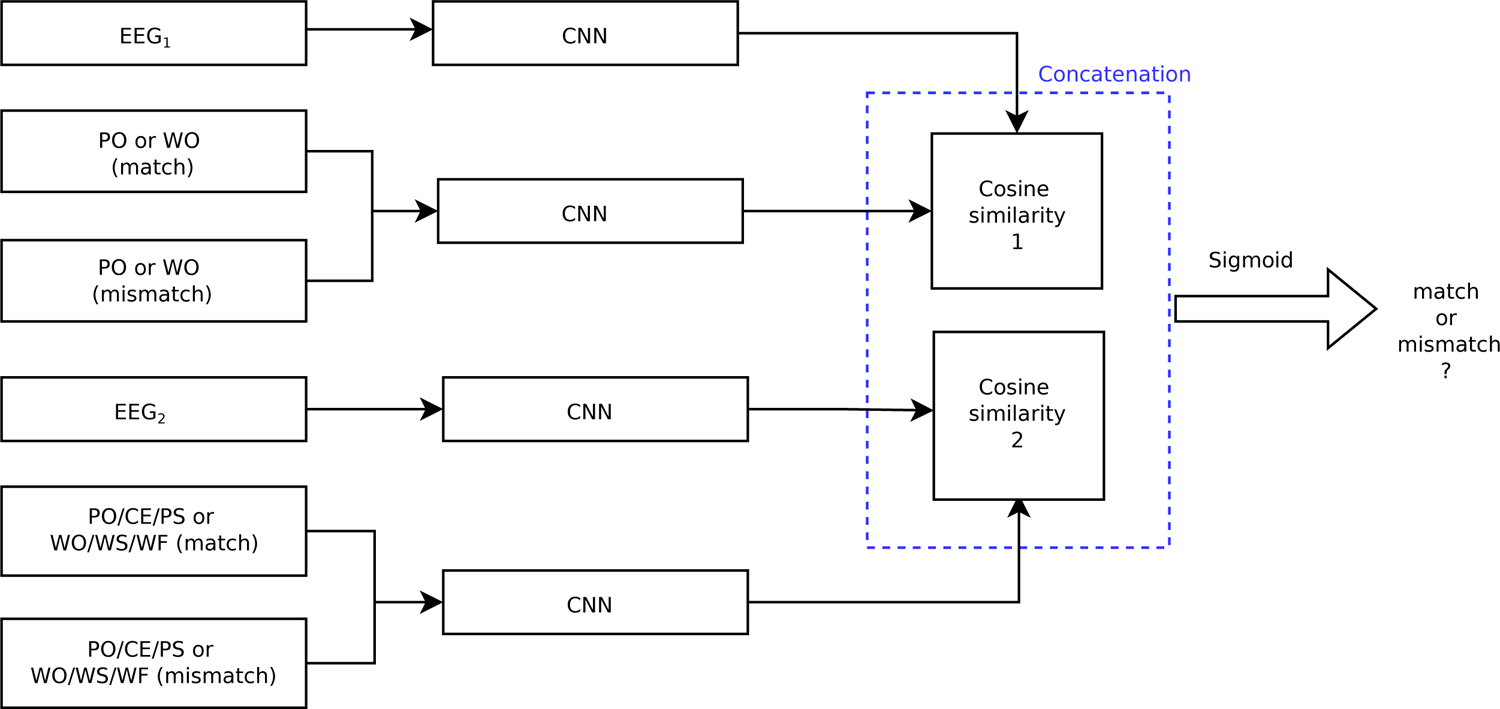
Linearized CNN/nonlinear CNN. This multi-input architecture allows the integration of two synchronized speech features and their corresponding EEG: *EEG*_1_ and *EEG*_2_, respectively. The possible phoneme-level features are PO, and either PO, CE, or PS, and the word-level features are WO and either WO, WS or WF. The EEG segment and their corresponding speech segments pass through different CNN blocks, and their embedding is further used to select which speech segment matches the EEG segment. *Robust neural tracking of linguistic speech representations using a convolutional neural network*12

As defined in the MM task, three inputs are provided to the model per speech feature (i.e., one EEG, one matched speech segment, and one mismatched speech segment). In the figure, there are two of these triplets, one for the lexical speech (*EEG*_1_, PO (match), PO(mismatch)), and one for the linguistic or second lexical speech feature (*EEG*_2_, PO/CE/PS (match), PO/CE/PS (mismatch)). Two separate EEG streams are created to constrain the model to extract EEG information solely related to the corresponding speech feature.

The EEG (size: 64 *× T*, 64 the number of EEG channels, and T the selected number of time samples) is passed through two convolutional blocks: one spatial convolution (kernel size of 1-time sample), and three temporal convolutions (each with a kernel size of 39 time samples or 609 ms) to model spatiotemporal information from the EEG input. For each convolution, a rectified linear unit (ReLU) function was used as activation.

The speech inputs (matched and mismatched) are processed in another stream with solely the three temporal convolutions. The speech input is always 1-dimensional as it has no spatial component to the model. Once the encoded representation of both EEG and speech segments are obtained, two normalized dot products (i.e., cosine similarity) are computed: one between EEG and the matched segment encodings, and one between EEG and the mismatched segment encodings, resulting in two 16 *×* 16 similarity matrices concatenated into a single 32 *×* 16 matrix. This operation is repeated for each speech feature, resulting in two 32 *×* 16 similarity matrices. These are concatenated into a 64 *×* 16 matrix that is flattened into a 1024 *×* 1 vector that is lastly passed through a dense layer with a sigmoid as the activation function. The model decides whether the first speech segment is a match or a mismatch based on the similarity between the encoded EEG and speech representations. Binary cross-entropy is used as the loss function on the match and mismatch labels.

#### 2.3.4. Linearized CNN

The linearized CNN is a variation of the nonlinear CNN, in which all ReLU activation functions of all the convolutions are replaced with linear functions to remove nonlinearities.

### 2.4. Model training

Every model can be trained according to three paradigms: subject-specific (SS), subject-independent (SI), or fine-tuned (FT). The SS paradigm involves using the training and validation sets (according to the split defined in Section 2.1.2) of a single subject to optimize the model and evaluate it on the unseen test set of the selected subject. The SI paradigm uses the combined training and validation sets of the 60 subjects to optimize the model and evaluate its performance per subject on their individual unseen testing set. Finally, the FT paradigm loads the weights obtained using the SI paradigm and fine-tunes the model on a selected subject’s training and validation sets before the evaluation on the selected subject’s test set.

### 2.5. Definition and role of lexical control and linguistics models

#### 2.5.1. Lexical control models

We use the lexical control models to ensure the linguistics models also use the linguistic information carried by each phoneme or word rather than just the timing of their onset. If the linguistic model performs significantly better than our control models, we can confirm the previous statement. In the control models, instead of providing two different speech features to the model, we provide the same feature twice, either twice PO (2PO) or twice WO (2WO). As a result, lexical models have the same number of parameters as the linguistics models while not incorporating linguistic information. We use 2PO as the control for phoneme-based linguistics models and 2WO for world-level linguistics models.

Although previous studies used acoustic controls, lexical controls with our current nonlinear CNN architecture might help us gain more insight into what the model uses in speech features.

Acoustics features are very broad and close to the original raw speech signal, and a possibility is that a powerful model could extract all the linguistic information from them, hence reducing the performance difference between the control and the linguistic models. We, therefore, choose more specific features as controls.

PO (or WO) is an impulse signal, having the value of 1 when the onset of a phoneme (or a word) occurs and 0 when it does not. The pulses are 1 sample long. Linguistic features are structurally the same (i.e., CE and PS are non-zero at the same time indices as PO; WS and WF are non-zero at the same time indices as WO) as depicted in Figure 1. However, the non-zero values have a varying magnitude that provides linguistic information. Therefore, if linguistics models outperform lexical control models, we can conclude that there is information in the EEG related to the onsets’ magnitude. We thus use the difference in performance (i.e., match-mismatch accuracy) between lexical control models and linguistics models as an objective measure of linguistic tracking.

#### 2.5.2. Linguistics models

The linguistics models we use in this study integrate both lexical and linguistic information. We investigate whether the model uses information contained in linguistics but not in lexical features to solve the MM task.

*Phoneme-based linguistics model*: We introduce two models including:

PO and CE (PO+CE)
PO and PS (PO+PS)

*Word-based linguistics model*: We introduce two models including:

WO and WS (WO+WS)
WO and WF (WO+WF)

2.6. *Statistical significance*

In Section 3.1, we compare performance across models using Wilcoxon signed-rank tests with an alpha level of 0.05, with Holm-Bonferroni corrections performed for word- and phoneme-level. All the tested models are reported in Table 1.

**Table 1:**
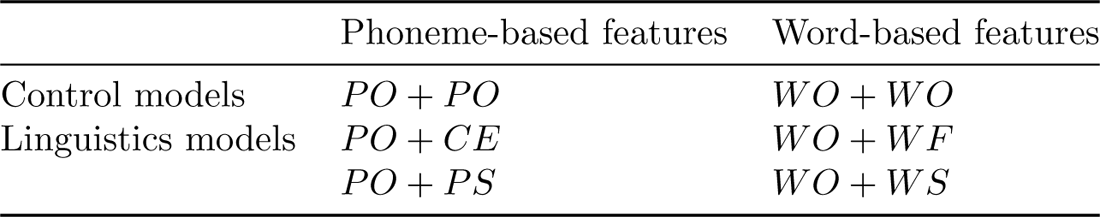
Overview of all the models evaluated. The control and linguistic models are reported as well as the feature combination they used. PO=phoneme onsets; PS=phoneme surprisal; CE=cohort entropy; WO=word onsets; WS=word surprisal; WF=word frequency.

We provide in Appendix B an analysis using linear mixed models (LMMs) to investigate the interactions between the different modalities of a model and its performance on the MM task. The modalities we investigated are the architecture, the training paradigm and the speech features provided to the model.

### 2.7. Data collection

We recorded the EEG of 60 subjects to which we presented 10 stories in Flemish, with each an average duration of 14 min 30 seconds (i.e., 145 min per subject). The EEG data from 18 of these 60 subjects were made publicly available, as part of a larger dataset (Bollens et al., 2023). More details about the experimental setup are reported in Appendix A.

## 3. Results

### 3.1. Neural tracking of linguistic features unrelated to onsets

#### 3.1.1. Phoneme-level linguistics

We here use the nonlinear CNN architecture, trained and validated on the training and validation sets of all the 60 subjects (subject-independent) and evaluated on each subject’s test set individually. Each point in the violin plots in Figure 5a is the accuracy obtained for individual subjects on their test sets. We compare the accuracy obtained on the MM task for each model on each subject’s test set between two phoneme-level linguistics models (PO+PS and PO+CE) and our phoneme-level lexical control model (2PO). On the right side of the figure, the paired difference in accuracy between each of the two PO+PS and PO+CE models and the 2PO model is depicted.

**Figure 5:**
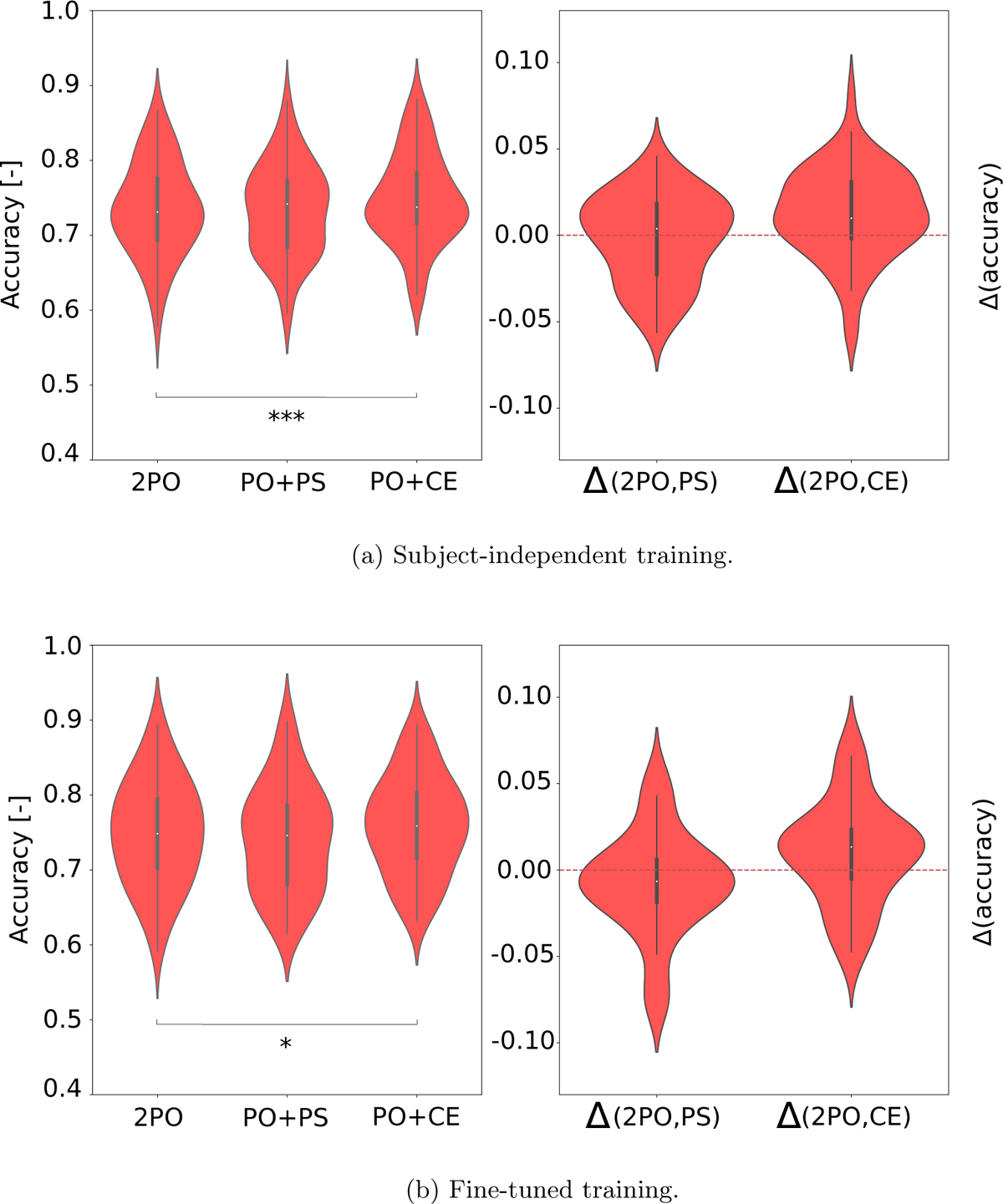
Phoneme-level linguistic tracking over phoneme onset for subject-independent and fine-tuned training paradigms using the nonlinear CNN architecture (window length: 5 s). The left panel depicts the MM accuracy obtained for each subject. The right panel depicts the difference in the MM accuracy between the linguistics and the control model (Δ(2*PO, PS*) and Δ(2*PO, CE*) respectively).(*∗*: *p <* 0.05*, ∗∗*: *p <* 0.01*, ∗ ∗ ∗*: *p <* 0.001)

We observe no significant difference between the linguistics model PO+PS and the lexical control model 2PO (*W* = 853, *p* = 0.812, after Holm-Bonferroni correction). We do observe a significant performance increase in the linguistics model PO+CE over the lexical control model 2PO (*W* = 410, *p <* 0.001, after Holm-Bonferroni correction).

We then repeat the same experiment while fine-tuning the model on each subject’s training and validation set before evaluating it on that subject’s test set. The results are depicted on Figure 5b.

We observe no significant difference between the linguistics model PO+PS and the lexical control model 2PO (*W* = 593, *p* = 0.0275, after Holm-Bonferroni correction). We do observe a significant performance increase of the linguistics model PO+CE over the lexical control model 2PO (*W* = 573, *p* = 0.0370 after Holm-Bonferronni correction).

#### 3.1.2. Word-level linguistics

The training and evaluation are performed identically to Section 3.1.1. We compare two word-level linguistics models (WO+WS and WO+WF) to our word-level lexical control model (2WO). The accuracy obtained on the MM task for each model on each subject’s test set is depicted in Figure 6a. On the right part of the figure, the paired difference in accuracy between each of the two WO+WS and WO+WF models and the 2WO model is depicted.

**Figure 6:**
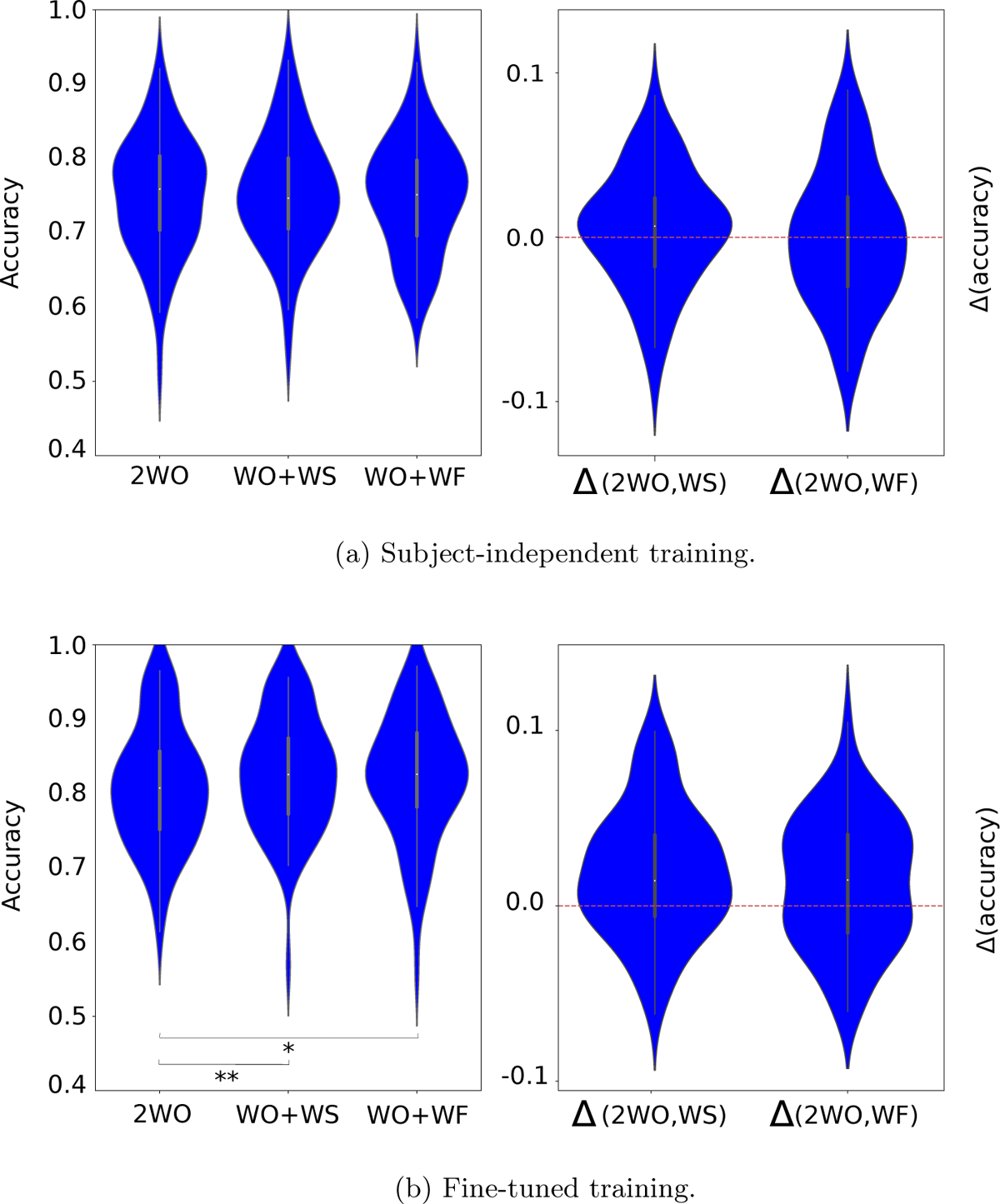
Word-level linguistic tracking over word onset for subject-independent and fine-tuned training paradigms using the nonlinear CNN architecture (window length: 10 s). The left panel depicts the MM accuracy obtained for all subjects on their test set using the lexical control model (2WO) and the linguistics models (WO+WS and WO+WF). The right panel depicts the difference in the MM accuracy between the linguistics and the control model (Δ(2*WO, WS*) and Δ(2*WO, WF*) respectively).(*∗*: *p <* 0.05*, ∗∗*: *p <* 0.01*, ∗ ∗ ∗*: *p <* 0.001) CNN-nonlinear CNN: *W* = 741, *p* = 0.277).

We observe no significant difference between linguistics models (WO+WS and WO+WF) and the lexical control model 2WO (*W* = 749, *p* = 0.444, and *W* = 841, *p* = 0.911 respectively, after Holm-Bonferroni corrections).

We then repeat the same experiment while fine-tuning the model on each subject’s training and validation set before evaluating it on each subject’s test set. The results are depicted on Figure 6b.

We observe a significant performance increase for the WO+WS model (*W* = 411, *p* = 0.00116, after Holm-Bonferroni correction) and the WO+WF (*W* = 570, *p* = 0.011, after Holm-Bonferonni correction) over the lexical control model 2WO.

### 3.2. Benefits of deep learning models to measure linguistic features tracking

The difference in accuracy between linguistics and lexical control models is indicative of neural tracking of linguistics, and the architecture significantly impacts the accuracy of models. We hence quantify the impact of the model’s architecture on this difference.

In Figure 7a, we depict the difference in accuracy between phoneme-based linguistics and their control models (Δ(*PO, PS*), and Δ(*PO, CE*)) for the linear encoder baseline, the linearized CNN and the nonlinear CNN. We report only the fine-tuned training condition as we observed the largest linguistics effect with it in subSection 3.1. The accuracy difference Δ(*PO, CE*) was significantly higher for the nonlinear CNN over the linear encoder baseline (*W* = 202, *p <* 0.001) and the linearized CNN (*W* = 249, *p <* 0.001). There was no significant difference between the linearized CNN and the linear encoder baseline (*W* = 769, *p* = 0.381). The accuracy difference Δ(*PO, PS*) did not change significantly across architectures (nonlinear CNN-linear SoA: *W* = 692, *p* = 0.145; linearized CNN-linear SoA: *W* = 798, *p* = 0.514; linearized

**Figure 7:**
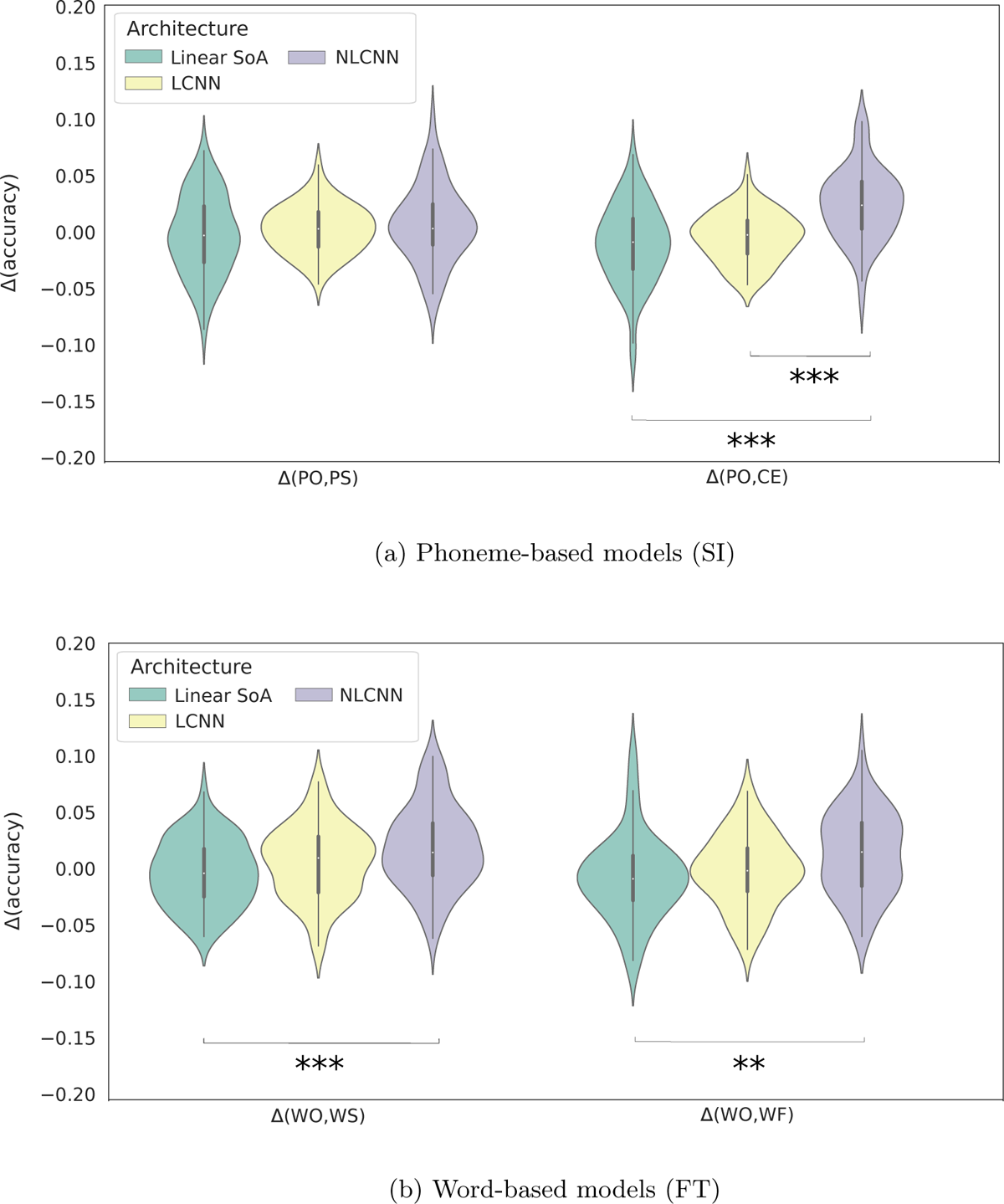
Difference in the MM accuracy between linguistic and lexical control models for phoneme-based (Figure a, 5 s segments) and word-based (Figure b, 10 s segments) models. Linear SoA=linear encoder baseline; NLCNN=nonlinear CNN; LCNN=linearized CNN. (*∗*: *p <* 0.05*, ∗∗*: *p <* 0.01*, ∗ ∗ ∗*: *p <* 0.001)

In Figure 7b, we depict the difference in accuracy between word-based linguistics and their control models (Δ(*WO, WS*), and Δ(*WO, WF*)) for the linear encoder baseline, the linearized CNN and the nonlinear CNN. We report only the subject-independent training condition as we solely observed a linguistics effect with it in subSection 3.1. The accuracy difference Δ(*WO, WS*) was significantly higher for the nonlinear CNN model than the linear encoder baseline (*W* = 393, *p <* 0.001), but not higher than the linearized CNN (*W* = 612, *p* = 0.0257, non-significant after Holm-Bonferroni correction). There was also no significant difference between the linearized CNN and the linear baseline encoder (*W* = 717, *p* = 0.146). The accuracy difference Δ(*WO, WF*) was significantly higher for the nonlinear CNN model over the linear encoder baseline (*W* = 500, *p* = 2.28 *∗* 10*^−^*^3^), but not over the linearized CNN (*W* = 599, *p* = 0.0202, non-significant after Holm-Bonferroni correction). There was also no significant difference between the linearized CNN and the linear baseline encoder (*W* = 710, *p* = 0.132).

## 4. Discussion

We first used our nonlinear CNN to evaluate if linguistic features significantly contribute beyond onset neural tracking for both phoneme- and word-based features.

At the phoneme level, and for both subject-independent and fine-tuned training conditions (see Section 3.1.1), cohort entropy added on top of phoneme onsets significantly contributes beyond phoneme onsets. Gillis et al. (2022); Brodbeck et al. (2018) showed the significant contribution of cohort entropy over acoustic and lexical features within a regression task; however they also showed the significant contribution of phoneme surprisal, which we did not observe. The use of different control methods, the complexity, and the non-linearity introduced by the nonlinear CNN could explain this. Gillis et al. (2022) evaluated neural tracking using a regression task as opposed to our match-mismatch task, which might also impact the conclusions.Brodbeck et al. (2018), the brain signals were recorded using magnetoencephalography (MEG), which might also explain effect differences.

At the word level, and only when fine-tuning, word surprisal and word frequency significantly contributed beyond word onsets (see Section 3.1.2). Gillis et al. (2022) showed the significant contribution of word surprisal and word frequency over and beyond acoustic and lexical features within a regression task; which is coherent with our current results. Possibly for the same reasons stated at the phoneme level, some results deviate from conclusions found in Gillis et al. (2022); Weissbart et al. (2019).

We conducted a last experiment in which we demonstrated that our nonlinear CNN shows a wider performance gap between linguistics models and their respective controls than the linear encoder baseline. This suggests that the nonlinearity and the complexity of a CNN model improves its ability to find the added value of linguistics tracking over onsets.

Although some studies argue in favor of acoustic features in the control models (Gillis et al. (2022); Brodbeck et al. (2018)), we here considered only a lexical control. An argument supporting our approach is that acoustic features (e.g., the mel spectrogram (Davis and Mermelstein, 1980)) contain some aspects of linguistic information (Gwilliams et al., 2022), and therefore including acoustic features in the control model, also removes part of the linguistic information from the final outcome measure, thereby reducing the power of the analysis. On the contrary, comparing a lexical and a linguistic feature (e.g., PO vs. PS) facilitates the interpretation: the unique difference between them is the pulse magnitude.

Cognitive effects, such as attention, could also impact our models’ performance as we did not explicitly model them here. For instance, a decrease in attention is often associated with a decrease in neural tracking of acoustic features (O’Sullivan et al., 2015; Ding and Simon, 2012). We do not consider attention processing in our model, so modulations of our models’ performance would be difficult to evaluate. To avoid such events, we required participants to perform an attention control task during recordings as described in Appendix A.

Apart from a study linearizing parts of deep learning architectures (Keshishian et al., 2020), the use of nonlinear models was mainly oriented towards performance and not interpretation. Our multi-input feature model, first introduced by Puffay et al. (2022), allows us to separate the processing of features with their corresponding EEG, which enables us to know what weight the model gives to each feature in various conditions (e.g., different SNRs). This will be covered in a future study dedicated to model interpretation.

## 5. Conclusion

Our study was conducted to measure the neural tracking of linguistic information that is unrelated to onset information and to investigate the benefits of DNNs over linear encoders to this end. We first compared the MM task performance of lexical control models with the linguistics model and found that the linguistics model performed significantly better under certain training conditions. In the second part, we quantified the difference in the match-mismatch accuracy between linguistics and control models across different architectures and confirmed the superiority of our nonlinear CNN over linear baselines.

## Acknowledgements

The authors thank all the participants for the recordings, as well as Wendy Verheijen, Kyara Cloes, Amelie Algoet, Jolien Smeulders, Lore Kerkhofs, Sara Peeters, Merel Dillen, Ilham Gamgami, Amber Verhoeven, Lies Bollens, Vitor Vasconcelos and Amber Aerts for their help with data collection. Funding was provided by the KU Leuven Special Research Fund C24/18/099 (C2 project to Tom Francart and Hugo Van hamme), FWO fellowships to Bernd Accou (1S89622N), Marlies Gillis (1SA0620N), Corentin Puffay (1S49823N), and Jonas Vanthornhout (1290821N).

## Appendix A. Data collection procedure

Sixty normal-hearing native Flemish speaking participants between 18 and 30 years old were recruited. This study was approved by the Medical Ethics Committee UZ KU Leuven/Research (KU Leuven, Belgium) with reference S57102, and all participants provided informed consent. We presented natural running speech (stories) in Flemish without background noise to participants and recorded the EEG signal simultaneously. All participants were normal hearing as confirmed by pure tone audiometry, and the Flemish Matrix test (Luts et al., 2014). All the participants listened to 10 unique stories of roughly the same duration (average: 14 minutes 30 seconds). The presentation order of the stories was randomized for each subject. The stories were presented binaurally at 62 dBA with shielded ER-3A insert phones (Etymotic, Elk Grove Village, Illinois, United States). Their only task was to answer a comprehension question after listening to each story to ensure they paid attention. EEG data were recorded using a 64-channel Active-Two EEG system (BioSemi, Amsterdam, The Netherlands) at a 8192 Hz sampling rate. The stimuli were presented using the APEX 4 software platform developed at ExpORL (Francart et al., 2008). The experiments took place in an electromagnetically shielded and soundproofed cabin.

## Appendix B. LMM analysis

### Appendix B.1. LMM models

We build a linear mixed model (LMM) given a series of predictors, and all their possible interactions using the R software package, and retained the best fit using, the Buildmer toolbox (Voeten, 2023). Separately for phoneme- and word-level models, we investigated the impact of linguistic features (L), training paradigm (Tr), and the model’s architecture (A) on the match-mismatch accuracy. We added subject ID (subject) as a random effect to the model. The full model before optimization is summarized in Equation B.1.

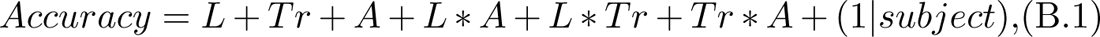

where L is a categorical variable taking as values either “PO”, “PS” or “CE” for phoneme-based models, and either “WO”, “WS or “WF for word-based models. Tr is a categorical variable with “SS”, “SI” or “FT” as possible values, and A indicates the architecture of a given model (“linear”, “linearized CNN” or “nonlinear CNN”).

The “*∗*” represents multiplication to model the interaction between variables. These variables are defined as fixed effects while subject is defined as a random effect. Using the Buildmer package from R, we use the maximum likelihood estimation procedure resulting in the model reported in Equation B.2 below.

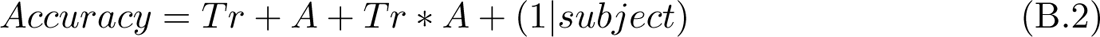

The models’ outcomes are reported in Section 3 for each source of variation, with the sum of squares, the mean squares, the F value and the corresponding p-value. If significant interaction effects were found or if we aimed to identify differences between values of a predictor, additional Holm-adjusted posthoc tests were performed on the estimated marginal means, linear trends or pairwise comparisons of these estimates, implemented by the Emmeans toolbox (Lenth et al., 2018).

### Appendix B.2. Effect of training paradigm, presence of linguistics and model architecture

#### Appendix B.2.1. Phoneme-level linguistic features

For each phoneme-based model, we depict in Figure B1a, the match-mismatch accuracy for the linear encoder baseline, the linearized CNN and the nonlinear CNN under subject-specific (SS), subject-independent (SI) and fine-tuned (FT) paradigms.

The LMM models described in Equation B.2 are then fitted and the outcomes of the ANOVA tests are depicted in Table B1. The presence of linguistics was not significant after model optimization (*F* = 1.14, *p* = 0.320) and hence was not reported. The training paradigm (*F* = 107, *p <* 0.001) and the architecture (*F* = 181 *∗* 10, *p <* 0.001) of the model have a significant impact. Finally, the interaction between the training paradigm and architecture was found to be significant (*F* = 83.6, *p <* 0.001).

Holm-adjusted pairwise comparisons confirmed that the average model accuracy over architecture levels was higher for the FT over the SI training paradigm (on average 0.0187 higher; *SE* = 0.00225, *df* = 1539, *t − ratio* = 8.30, *p <* 0.001) and for the SI over SS training paradigm (on average 0.0142 higher; *SE* = 0.00225, *df* = 1549, *t − ratio* = 6.30, *p <* 0.001).

As the interaction term *Tr ∗ A* is significant, we provide more details about the average trends stated above. The FT training paradigm generally performed significantly better than the SS training paradigm (*t − ratio* = 19.5, *p <* 0.001 for the nonlinear CNN, and *t − ratio* = 5.75, *p <* 0.001 for the linearized CNN), except for the linear encoder baseline, where no difference was observed (*t − ratio* = 0.047, *p* = 1.0). On the other hand, the effect of FT over SI was not consistent across architectures, for the nonlinear CNN, the difference was not significant (*t − ratio* = 2.93, *p* = 0.0823), whereas it was significant for both the linearized CNN (*t − ratio* = 4.13, *p* = 0.0013) and the linear encoder baseline (*t − ratio* = 7.3, *p <* 0.001). Finally, SI outperformed SS for the nonlinear CNN (*t − ratio* = 36.8, *p <* 0.001), but not for the linearized CNN (*t − ratio* = 1.62, *p* = 0.794). With the linear encoder baseline, SS outperformed SI (*t − ratio* = *−*7.25, *p <* 0.001).

### Appendix B.2.2. Word-level linguistics

For each word-based model, we depict in Figure B1b, the match-mismatch accuracy for the linear encoder baseline, the linearized CNN, and the nonlinear CNN under subject-specific (SS), subject-independent (SI) and fine-tuned (FT) paradigms.

The LMM models are then evaluated and the outcomes of the ANOVA tests are depicted in Table B2. The presence of linguistics was not significant after model optimization (*F* = 0.0956, *p* = 0.901), hence not reported. The training paradigm (*F* = 258, *p <* 0.001) and the architecture (*F* = 160 *∗* 10, *p <* 0.001) of the model have a significant impact. Finally, the interaction between the training paradigm and architecture was found to be significant (*F* = 104, *p <* 0.001).

A Holm-adjusted pairwise comparison confirmed that the average model accuracy over training paradigm levels was higher for the nonlinear CNN over the linearized CNN (on average 0.125 higher; *SE* = 0.00307, *df* = 155 *∗* 10, *t − ratio* = 40.7, *p <* 0.001) and for the linearized CNN over the linear encoder baseline (on average 0.0421 higher; *SE* = 0.00307, *df* = 1549, *t − ratio* = 13.7, *p <* 0.001).

Holm-adjusted pairwise comparisons confirmed that the average model accuracy over architecture levels was higher for the FT over the SI training paradigm (on average 0.0433 higher; *SE* = 0.00307, *df* = 1549, *t − ratio* = 14.0, *p <* 0.001) and for the SI over SS training paradigm (on average 0.0257 higher; *SE* = 0.00307, *df* = 1549, *t − ratio* = 8.37, *p <* 0.001).

As the interaction term *Tr ∗ A* is significant, we provide more specific details about the average trends above. The FT training paradigm was overall performing significantly better than the SS training paradigm (*t − ratio* = 28.9, *p <* 0.001, for the nonlinear CNN, and *t − ratio* = 3.44, *p* = 0.0047 for the linearized CNN) except for the linear encoder baseline, where no difference was observed (*t − ratio* = 0.067, *p* = 1.0). As opposed to phoneme-level models, FT consistently outperformed SI (*t − ratio* = 16.2, *p <* 0.001 for the nonlinear CNN, *t − ratio* = 3.42, *p* = 0.0047 for the linearized CNN, and *t − ratio* = 8.33, *p <* 0.001 for the linear encoder baseline). Finally, SI outperformed SS for the nonlinear CNN (*t − ratio* = 36.8, *p <* 0.001) and the linear encoder baseline (*t − ratio* = 27.6, *p <* 0.001), but not for the linearized CNN (*t − ratio* = 0.0022, *p* = 1.0).

---

**Figure B1:**
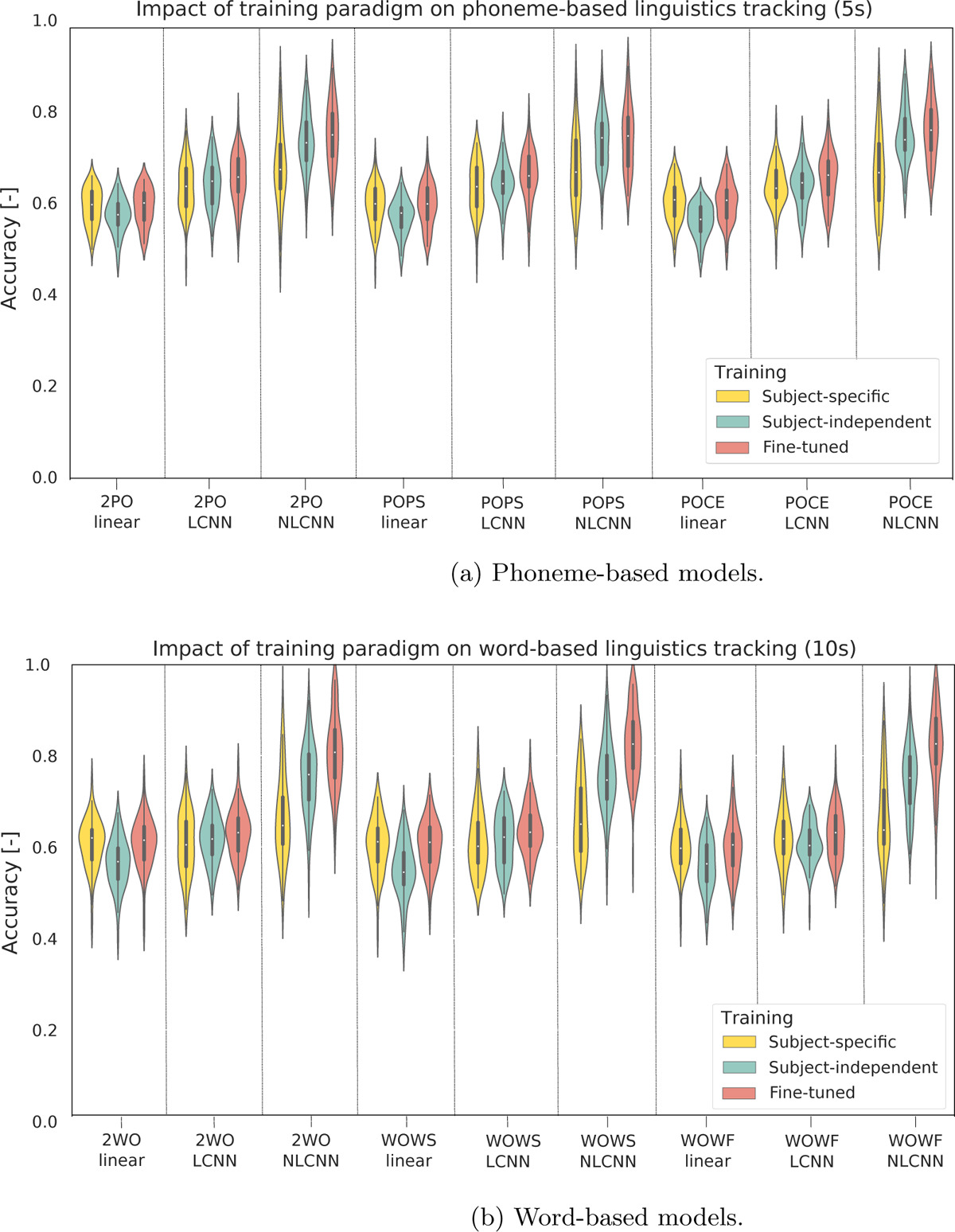
Impact of training conditions on word- and phoneme-based models. In the upper panel, the phoneme-based models are depicted (2PO, PO+PS, and PO+CE, respectively). In the lower panel, the word-based models are depicted (2WO, WO+WS, and WO+WF, respectively). For each feature combination, the linear encoder baseline (linear), the linearized CNN (LCNN), and the nonlinear CNN (NLCNN) models were trained in subject-specific (SS) and subject-independent (SI) paradigms. The linearized CNN and CNN models were also fine-tuned per subject (FT).

**Table B1:** ANOVA test results for phoneme-based models. Training=categorical variable with value “SS”, “SI” and “FT”; Architecture=categorical variable with value “linear”, “linearized CNN” or “CNN”. Holm-adjusted pairwise comparisons confirmed that the average model accuracy over training paradigm levels was higher for the nonlinear CNN over the linearized CNN (on average 0.076 higher; *SE* = 0.002, *df* = 1549, *t − ratio* = 34.2, *p <* 0.001) and for the linearized CNN over the linear encoder baseline (on average 0.058 higher; *SE* = 0.002, *df* = 1549, *t − ratio* = 25.9, *p <* 0.001).

| | Sum Square | Mean Square | F value | $Pr(< F)$ |
| --- | --- | --- | --- | --- |
| Training | 0.293 | 0.146 | 107 | $p < 0.001$ |
| Architecture | 4.93 | 2.46 | $181 \cdot 10^1$ | $p < 0.001$ |
| Training:Architecture | 0.455 | 0.113 | 83.6 | $p < 0.001$ |

**Table B2:**
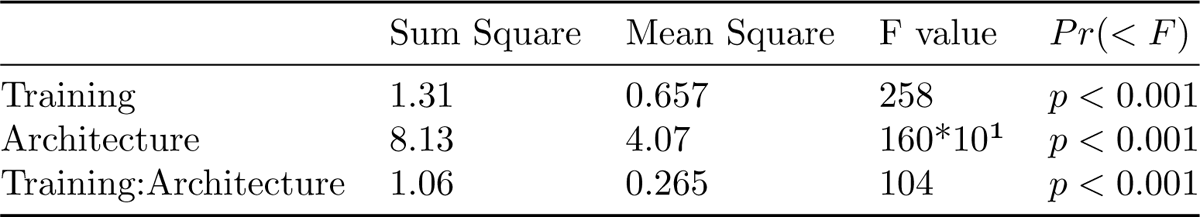
ANOVA test results for word-based models. Training=categorical variable with value “SS”, “SI” and “FT”; Architecture=categorical variable with value “linear”, “linear CNN” or “CNN”.

